# Bile acids and bilirubin effects on osteoblastic gene profile. Implications in the pathogenesis of osteoporosis in liver diseases

**DOI:** 10.1101/705871

**Authors:** Silvia Ruiz-Gaspà, Nuria Guañabens, Susana Jurado González, Marta Dubreuil, Andres Combalia, Pilar Peris, Ana Monegal, Albert Parés

## Abstract

Osteoporosis in advanced cholestatic and end-stage liver disease is related to low bone formation. Previous studies have demonstrated the deleterious consequences of lithocholic acid (LCA) and bilirubin on osteoblastic cells. These effects are partially or completely neutralized by ursodeoxycholic acid (UDCA). We have assessed the differential gene expression of osteoblastic cells under different culture conditions. The experiments were performed in human osteosarcoma cells (Saos-2) cultured with LCA 10 μM), bilirubin (50 μM) or UDCA (10 and 100 μM) at 2 and 24 hours. Expression of 87 genes related to bone metabolism and other signalling pathways were assessed by TaqMan micro fluidic cards. Several genes were up-regulated by LCA, most of them pro-apoptotic (*BAX*, *BCL10*, *BCL2L13*, *BCL2L14*), but also *MGP* (matrix Gla protein), B*GLAP* (osteocalcin), *SPP1* (osteopontin) and *CYP24A1*, and down-regulated bone morphogenic protein genes (*BMP3* and *BMP4*) and *DKK1* (Dickkopf-related protein 1). Parallel effects were observed with bilirubin, which up-regulated apoptotic genes and *CSF2* (colony-stimulating factor 2) and down-regulated antiapoptotic genes (*BCL2* and *BCL2L1)*, *BMP3*, *BMP4* and RUNX2. UDCA 100 μM had specific consequences since differential expression was observed, up-regulating *BMP2*, *BMP4*, *BMP7*, *CALCR* (calcitonin receptor), *SPOCK3* (osteonectin), *BGLAP* (osteocalcin) and *SPP1* (osteopontin), and down-regulating pro-apoptotic genes. Furthermore, most of the differential expression changes induced by both LCA and bilirubin were partially or completely neutralized by UDCA. *Conclusion*: Our observations reveal novel target genes, whose regulation by retained substances of cholestasis may provide additional insights into the pathogenesis of osteoporosis in cholestatic and end-stage liver diseases.

## Introduction

Osteoporosis is a skeletal disease characterized by low bone mass and micro-architectural deterioration of bone tissue, leading to an increased fragility and susceptibility to fracture. It is a common complication of liver diseases, particularly in chronic cholestasis and especially in those patients with primary biliary cholangitis (PBC) [1–4]. Low bone formation as a consequence of a deficient osteoblast activity is the main cause for bone loss [1], but an increased resorption has been described as well [5].

Different studies have found that high concentrations of bilirubin and bile acid can contribute to the abnormal osteoblast function [6], as both bilirubin and lithocholic acid (LCA) in addition to serum from jaundiced patients have detrimental effects on these bone-forming cells [7].

Ursodeoxycholic acid (UDCA), the standard treatment for patients with PBC has greatly changed the natural history of the disease [8–11]. Moreover, in bone cells UDCA increases survival and improves differentiation of human osteoblasts, neutralizing the detrimental effects of LCA and bilirubin on osteoblast survival, differentiation and mineralization [12]. Lastly, while bilirubin and LCA act as pro-apoptotic agents in human osteoblasts, UDCA has anti-apoptotic effects and neutralizes the apoptosis induced by LCA and bilirubin in osteoblastic cells [12].

In previous studies a down-regulation of RUNX2 gene by bilirubin and an up-regulation of the RANKL/OPG expression ratio by jaundiced sera in osteoblastic cells were demonstrated [7]. However, more data on the influence of the retained substances of cholestasis in osteoblastic cell gene expression is lacking. To gain new insights into cholestatic-induced osteoporosis, we have assessed whether the damaging effects of the retained substances such as bilirubin and bile acids can modify the gene expression profiling of osteoblastic cells.

## Materials and methods

### Materials

Dubelcco’s modified Eagle medium (DMEM), fetal bovine serum (FBS), Hanks Balanced Salt Solution (HBSS), L-glutamine and trypsin were purchased from Invitrogen (Grand Island, NY, USA); LCA, UDCA and bilirubin were from Sigma Chemical Co. (St. Louis, MO, USA); Penicillin-streptomicin was from LabClinics (Barcelona, Spain).

### Cell culture and incubation

The experiments were performed with human osteosarcoma cell line Saos-2. The cell line Saos-2 was obtained from the American Type Culture Collection (ATCC) (Rockville, MD, USA) (HTB85; ATCC) and cultured as a monolayer in DMEM containing 10% FCS, 100 U/mL penicillin and 100 mg/mL streptomycin. Cells were incubated at 37 °C in a humidified atmosphere of 5% CO^2^ in air.

### Administrated treatments

Saos-2 cells were cultured for 2 and 24 h in different conditions: (a) LCA (10 μM), bilirubin (50 μM) and UDCA 10 μM and 100 μM. (b) To analyze the interaction of UDCA with LCA and bilirubin, cells were incubated with a steady concentration of LCA (10 μM) or bilirubin (50 μM) and two concentrations of UDCA (10 μM and 100 μM).

All treatment concentrations used were selected based on the cytotoxicity assays carried out in previous studies [7,13].

### Experimental bilirubin solution preparation

Bilirubin (Sigma) stock solution of 1600 μM was prepared just before use by dissolving bilirubin in 10 ml 0.01N NaOH under dim light as previously described [12,15–18]. It was filtered through a sterile filter (0.22 μm pore size) and adjusted to pH 7.2-7.4 with 0.1N HCl, if necessary. The bilirubin stock solution was added to a final concentration of 50 μM in the culture medium. The cell cultures were kept in dark conditions to prevent bilirubin light degradation. Control cells were treated with vehicle (NaOH 0.1N).

### RNA isolation and quantification

Total cellular RNA was extracted from cultured cells using an acid guanidinium-phenol-chloroform method (Trizol reagent; Invitrogen, Grand Island, NY, USA) according to the manufacturer’s protocols. RNA integrity was determined by a microfluidics-based electrophoresis system using a 2100 Bioanalyzer (Agilent Techologies, Palo Alto, CA). RNA integrity number (RIN) from automated analysis software allows classification of RNA in a numeric system with one for complete degradation and ten for optimal intactness. Both analyses displayed highly intact RNA, with RIN values of 9.8–10.

### Expression analysis with TaqMan microfluidic cards

cDNA synthesis was performed with the High-Capacity cDNA Reverse Transcription Kit (Applied Biosystems, Foster City, CA, USA) with a Master Mix containing 2.5 U/μl of MultiScribe Reverse Transcriptase and 1 μg of total RNA. The reaction mixture was incubated at 25°C for 10 min, followed by 120 min at 37°C and then by heat inactivation of the enzyme at 85°C for 5 sec. Next, we mixed 2 μl of single-stranded cDNA (equivalent to around 100 ng of total RNA) with 48 μl of nuclease-free water and 50 μl of TaqMan Universal PCR Master Mix. After loading 100 μl of the sample-specific PCR mixture into one sample port of the microfluidic cards (Human ABC Transporter Panel; Applied Biosystems), the cards were centrifuged twice for 1 min at 280g and sealed to prevent well-to-well contamination. The cards were placed in the microfluidic card Sample Block of an ABI Prism 7900 HT Sequence Detection System (Applied Biosystems). The thermal cycling conditions were 2 min at 50°C and 10 min at 95°C, followed by 40 cycles of 30 sec at 97°C and 1 min at 59.7°C. The assay for each gene on the microfluidic card was carried out in triplicate, due to the design of this specific panel. The calculation of the threshold cycle (Ct) values were performed using the SDS 2.2 software (Applied Biosystems), after automatically setting the baseline and the threshold.

### Gene selection

A total of eighty-seven genes were selected to investigate the profile gene expression under bilirubin, LCA and UDCA treatment. Genes were chosen according to their relevant function in cellular processes and signaling pathways related to bone metabolism (Table 1).

**Table 1.**
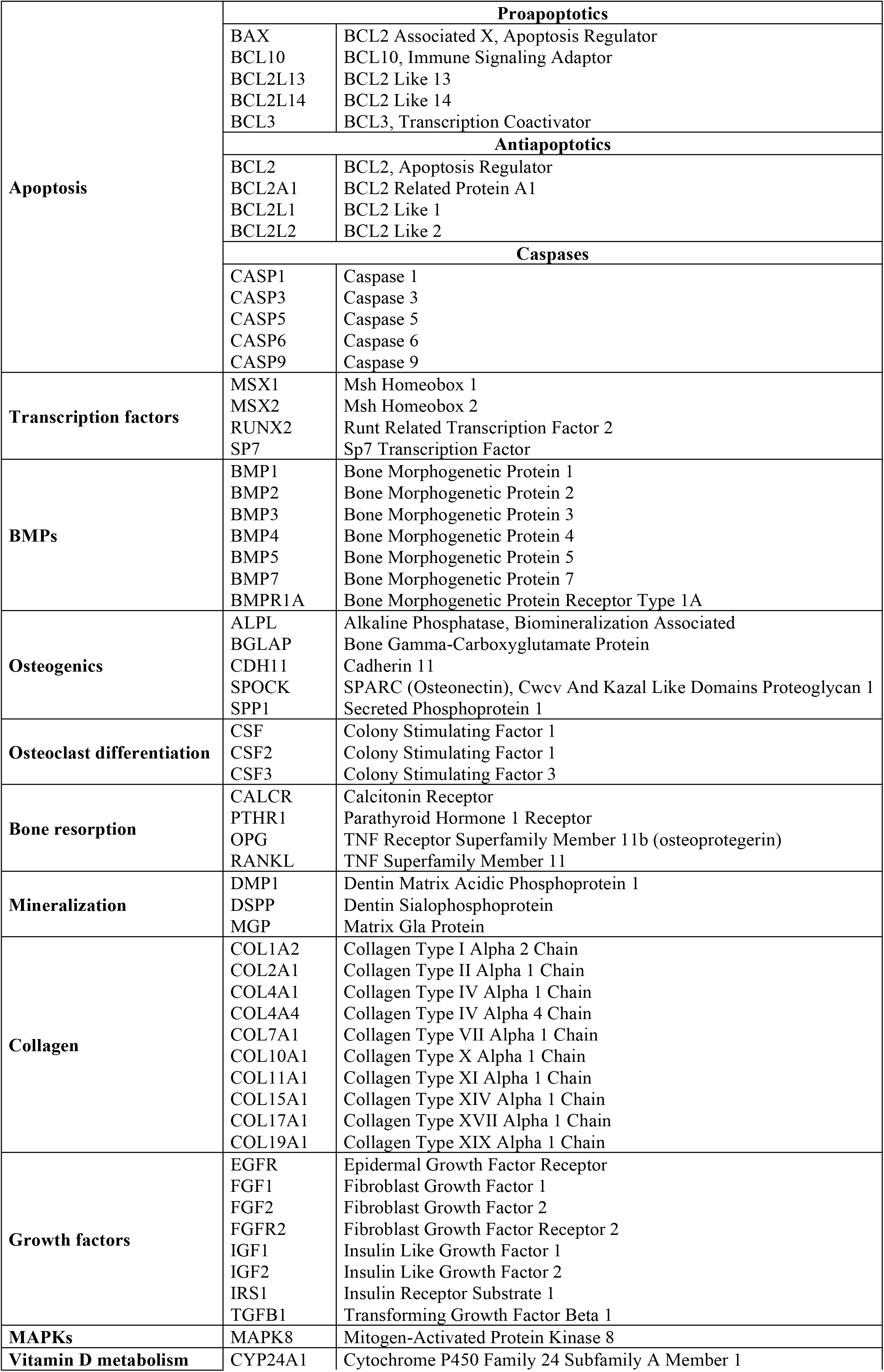
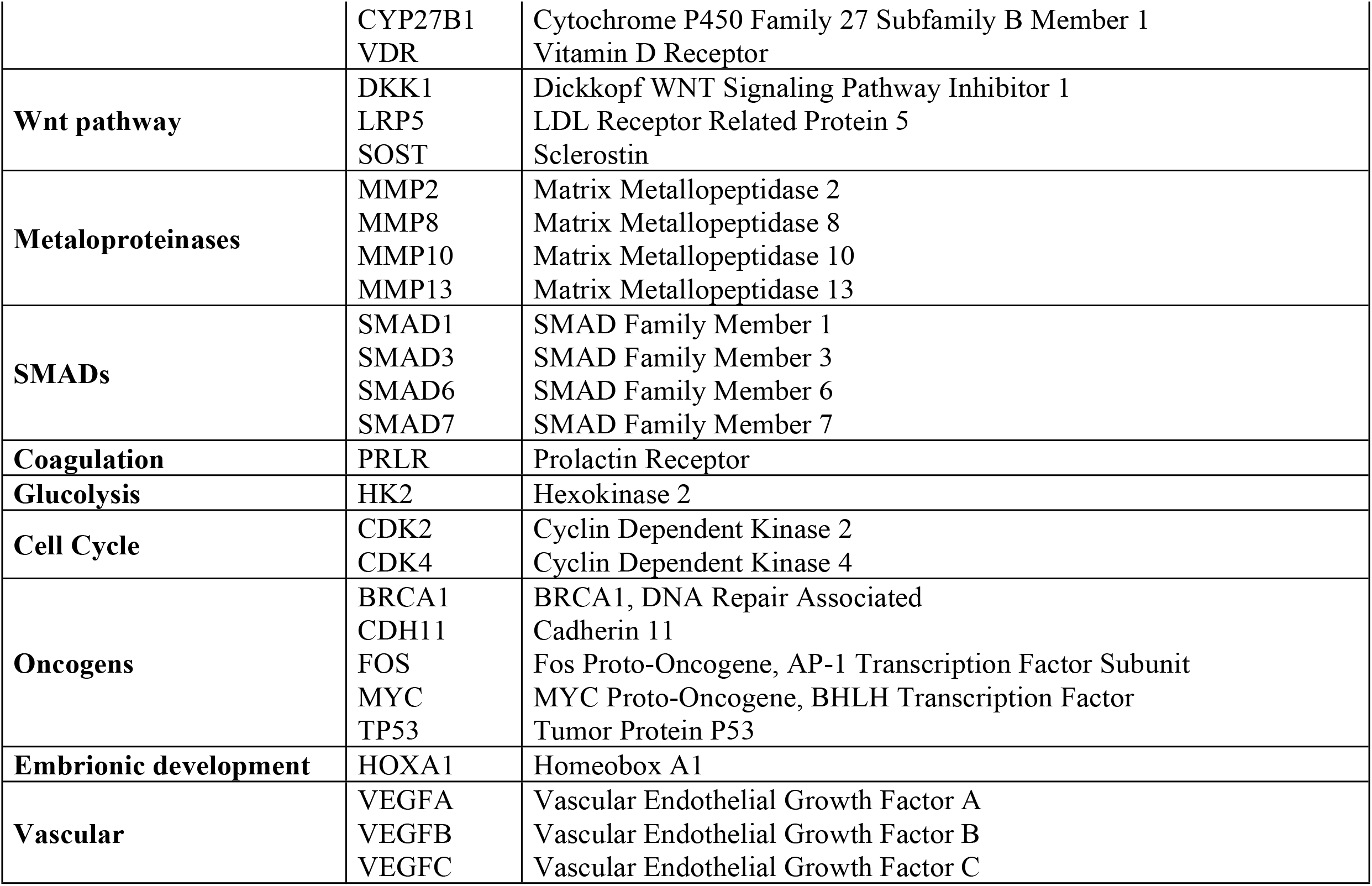
Selected genes.

|  |  |  |
| --- | --- | --- |
| **Apoptosis** | **Proapoptotics** |  |
|  | BAX<br>BCL10<br>BCL2L13<br>BCL2L14<br>BCL3 | BCL2 Associated X, Apoptosis Regulator<br>BCL10, Immune Signaling Adaptor<br>BCL2 Like 13<br>BCL2 Like 14<br>BCL3, Transcription Coactivator |
|  | **Antiapoptotics** |  |
|  | BCL2<br>BCL2A1<br>BCL2L1<br>BCL2L2 | BCL2, Apoptosis Regulator<br>BCL2 Related Protein A1<br>BCL2 Like 1<br>BCL2 Like 2 |
|  | **Caspases** |  |
|  | CASP1<br>CASP3<br>CASP5<br>CASP6<br>CASP9 | Caspase 1<br>Caspase 3<br>Caspase 5<br>Caspase 6<br>Caspase 9 |
| **Transcription factors** | MSX1<br>MSX2<br>RUNX2<br>SP7 | Msh Homeobox 1<br>Msh Homeobox 2<br>Runt Related Transcription Factor 2<br>Sp7 Transcription Factor |
| **BMPs** | BMP1<br>BMP2<br>BMP3<br>BMP4<br>BMP5<br>BMP7<br>BMPR1A | Bone Morphogenetic Protein 1<br>Bone Morphogenetic Protein 2<br>Bone Morphogenetic Protein 3<br>Bone Morphogenetic Protein 4<br>Bone Morphogenetic Protein 5<br>Bone Morphogenetic Protein 7<br>Bone Morphogenetic Protein Receptor Type 1A |
| **Osteogenics** | ALPL<br>BGLAP<br>CDH11<br>SPOCK<br>SPP1 | Alkaline Phosphatase, Biomineralization Associated<br>Bone Gamma-Carboxyglutamate Protein<br>Cadherin 11<br>SPARC (Osteonectin), Cwcv And Kazal Like Domains Proteoglycan 1<br>Secreted Phosphoprotein 1 |
| **Osteoclast differentiation** | CSF<br>CSF2<br>CSF3 | Colony Stimulating Factor 1<br>Colony Stimulating Factor 1<br>Colony Stimulating Factor 3 |
| **Bone resorption** | CALCR<br>PTH1R<br>OPG<br>RANKL | Calcitonin Receptor<br>Parathyroid Hormone 1 Receptor<br>TNF Receptor Superfamily Member 11b (osteoprotegerin)<br>TNF Superfamily Member 11 |
| **Mineralization** | DMP1<br>DSPP<br>MGP | Dentin Matrix Acidic Phosphoprotein 1<br>Dentin Sialophosphoprotein<br>Matrix Gla Protein |
| **Collagen** | COL1A2<br>COL2A1<br>COL4A1<br>COL4A4<br>COL7A1<br>COL10A1<br>COL11A1<br>COL15A1<br>COL17A1<br>COL19A1 | Collagen Type I Alpha 2 Chain<br>Collagen Type II Alpha 1 Chain<br>Collagen Type IV Alpha 1 Chain<br>Collagen Type IV Alpha 4 Chain<br>Collagen Type VII Alpha 1 Chain<br>Collagen Type X Alpha 1 Chain<br>Collagen Type XI Alpha 1 Chain<br>Collagen Type XIV Alpha 1 Chain<br>Collagen Type XVII Alpha 1 Chain<br>Collagen Type XIX Alpha 1 Chain |
| **Growth factors** | EGFR<br>FGF1<br>FGF2<br>FGFR2<br>IGF1<br>IGF2<br>IRS1<br>TGFB1 | Epidermal Growth Factor Receptor<br>Fibroblast Growth Factor 1<br>Fibroblast Growth Factor 2<br>Fibroblast Growth Factor Receptor 2<br>Insulin Like Growth Factor 1<br>Insulin Like Growth Factor 2<br>Insulin Receptor Substrate 1<br>Transforming Growth Factor Beta 1 |
| **MAPKs** | MAPK8 | Mitogen-Activated Protein Kinase 8 |
| **Vitamin D metabolism** | CYP24A1 | Cytochrome P450 Family 24 Subfamily A Member 1 |
|  | CYP27B1<br>VDR | Cytochrome P450 Family 27 Subfamily B Member 1<br>Vitamin D Receptor |
| **Wnt pathway** | DKK1<br>LRP5<br>SOST | Dickkopf WNT Signaling Pathway Inhibitor 1<br>LDL Receptor Related Protein 5<br>Sclerostin |
| **Metalloproteinases** | MMP2<br>MMP8<br>MMP10<br>MMP13 | Matrix Metalloproteinase 2<br>Matrix Metalloproteinase 8<br>Matrix Metalloproteinase 10<br>Matrix Metalloproteinase 13 |
| **SMADs** | SMAD1<br>SMAD3<br>SMAD6<br>SMAD7 | SMAD Family Member 1<br>SMAD Family Member 3<br>SMAD Family Member 6<br>SMAD Family Member 7 |
| **Coagulation** | PRLR | Prolactin Receptor |
| **Glucolysis** | HK2 | Hexokinase 2 |
| **Cell Cycle** | CDK2<br>CDK4 | Cyclin Dependent Kinase 2<br>Cyclin Dependent Kinase 4 |
| **Oncogens** | BRCA1<br>CDH11<br>FOS<br>MYC<br>TP53 | BRCA1, DNA Repair Associated<br>Cadherin 11<br>Fos Proto-Oncogene, AP-1 Transcription Factor Subunit<br>MYC Proto-Oncogene, BHLH Transcription Factor<br>Tumor Protein P53 |
| **Embryonic development** | HOXA1 | Homeobox A1 |
| **Vascular** | VEGFA<br>VEGFB<br>VEGFC | Vascular Endothelial Growth Factor A<br>Vascular Endothelial Growth Factor B<br>Vascular Endothelial Growth Factor C |

### Statistics

Significant differences between two groups were determined by Student’s t-test or Mann–Whitney U-test. When multiple groups were compared, ANOVA was utilized, followed by a Tukey’s multiple contrast test, when applicable. A p-value ≤ 0.05 was considered significant. All analyses were performed using the PASW Statistics 20 (SPSS, Chicago, IL, USA).

## Results

### Effect of LCA and bilirubin on gene profiles

As compared with controls the effects of LCA at 10μM and bilirubin at 50μM were observed after 2 hours of treatment, but frequently were more apparent after 24 hours. The most relevant results are shown in figures 1A and 1B.

**Fig 1.**
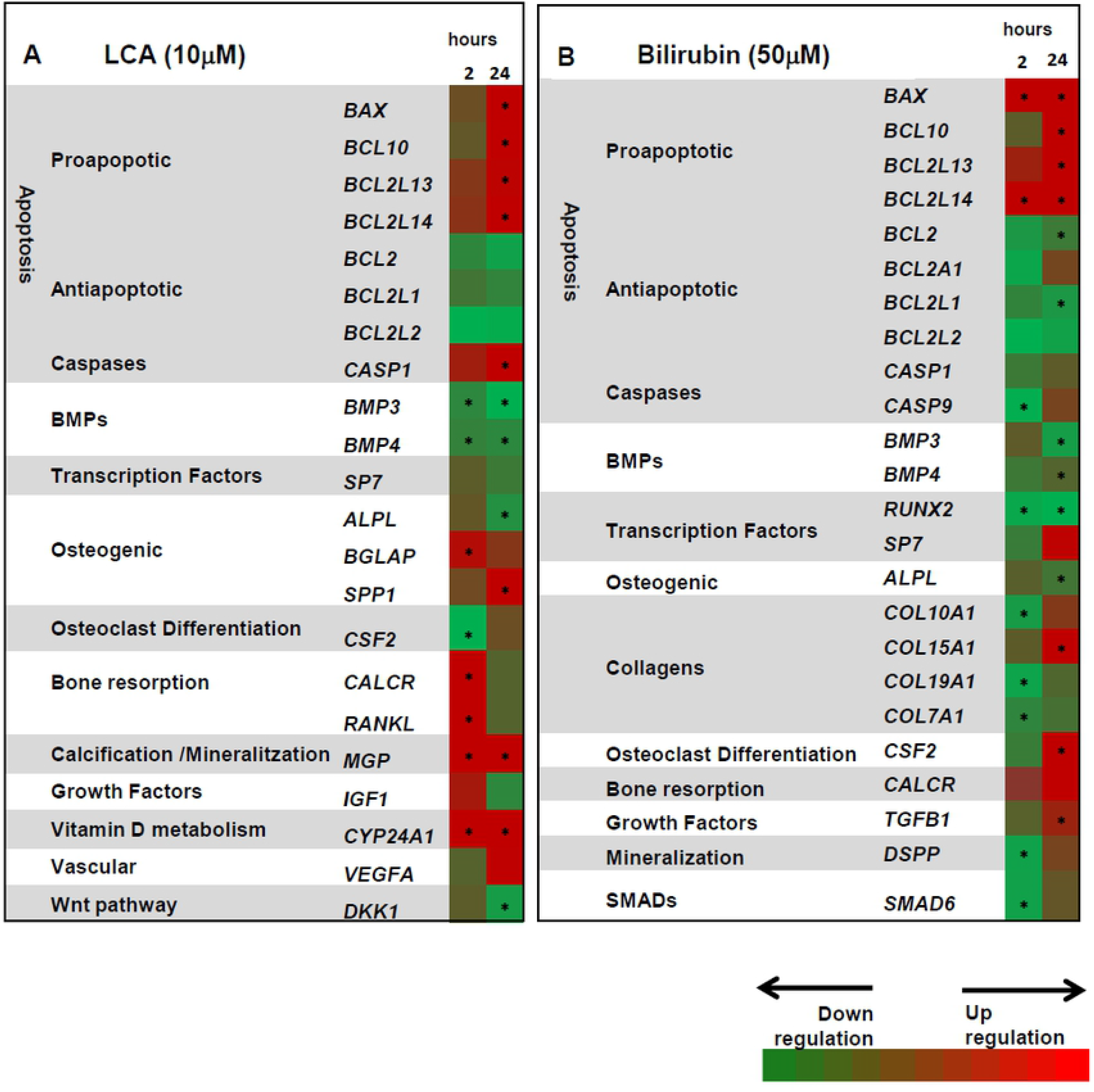
Heat map of the effects of litocholic acid (A) and bilirubin (B) on gene expression profiles in Saos-2 osteoblastic cells. The red colour corresponds to genes that are up-regulated, and green colour corresponds to genes that are down-regulated as compared to controls without LCA or bilirubin. * indicates significant differences vs non-treated cells.

The apoptosis-related genes were the ones most affected after treatments with LCA and bilirubin, resulting in severe gene expression changes. LCA and bilirubin significantly up-regulated the expression of some pro-apoptotic genes and down-regulated some anti-apoptotic genes (p < 0.05). Caspase 1 (*CASP1*) was overexpressed under LCA treatment and bilirubin decreased the expression of caspase 9 (CASP9).

When assessing bone morphogenetic proteins (BMPs), both LCA and bilirubin diminished the expression of some of these genes, mainly *BMP3* and *BMP4* (Figures 1A and 1B). These effects were observed after just two hours of treatment and were more evident under LCA treatment (figure 2).

**Fig 2.**
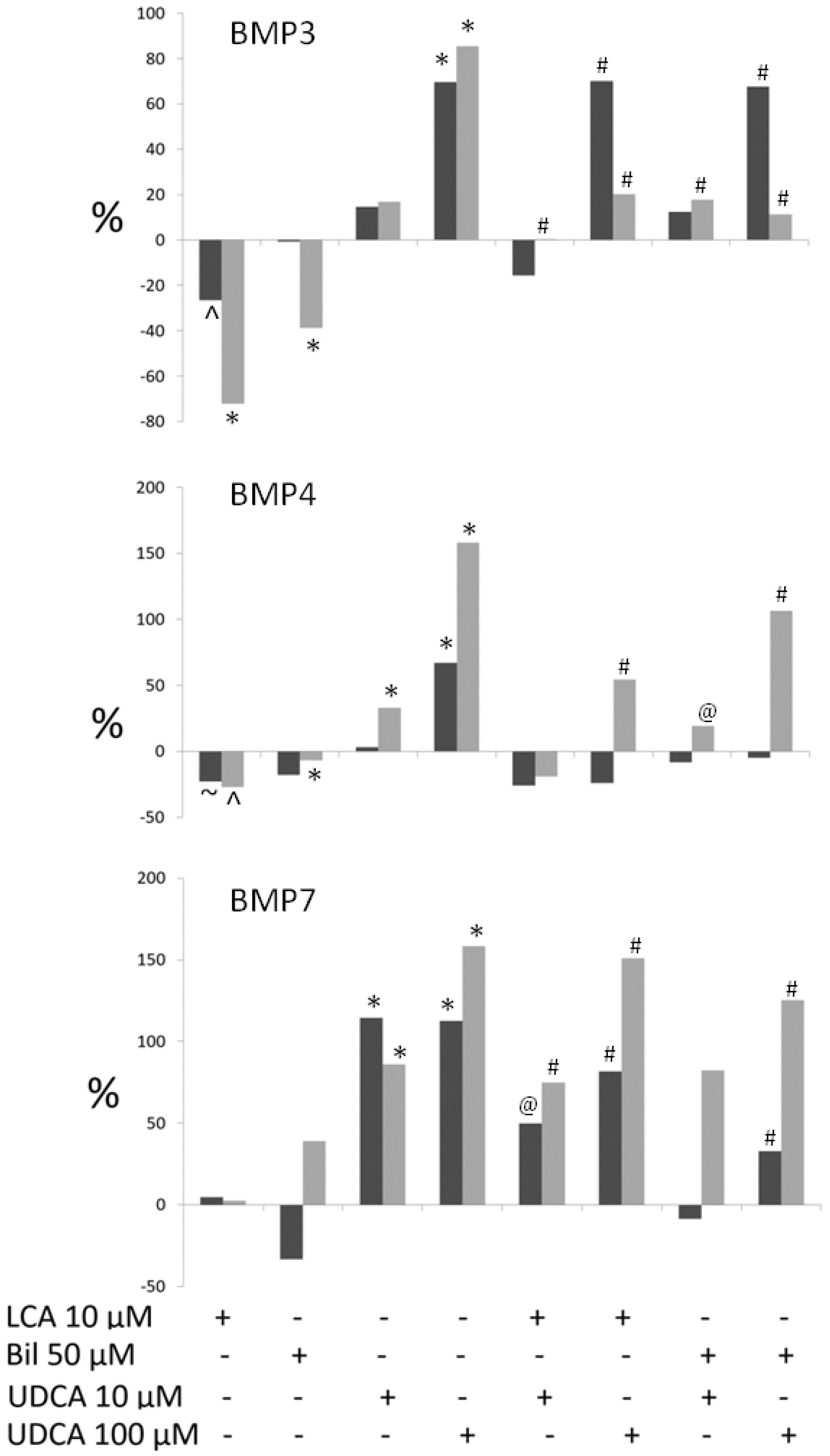
Percentage changes in bone morphogenetic protein (BMP3, BMP4 and BMP7) gene expression at 2 hours (dark grey bars) and 24 hours (light grey bars), in different culture conditions (LCA, Bilirubin, and UDCA) and the effect of UDCA on cells treated with LCA or bilirubin. (*p<0.001 vs controls; ^p<0.01 vs controls; ~p<0.05 vs controls; # p<0.001 vs UDCA treated cells; @ p<0.01 vs UDCA treated cells).

The expression of collagen X alpha-1 (*COL10A1*), collagen XIX alpha-1 (*COL19A1*) and collagen VII alpha-1 (*COL7A1*) was down-regulated by bilirubin after 2 hours of treatment. However, collagen XV alpha-1 (*COL15A1*) was up-regulated after 24 hours. No significant changes were observed with LCA on the different evaluated collagen genes.

The analysis of selected osteogenic genes displayed substantial overexpression of osteocalcin or bone gamma-carboxyglutamic acid-containing protein (*BGLAP)* and osteopontin (SPP1) (p<0.001) under LCA treatment. Moreover, a significant decrease of alkaline phosphatase expression *(ALPL)* was found under LCA treatment (p<0.001) and under bilirubin treatment at 24 hours (p<0.05).

Regarding the specific transcription factors, SP7 (osterix transcription factor) expression was down-regulated after 24 hours under LCA treatment, while it was up-regulated after 24 hours with bilirubin treatment. The runt-related transcription factor 2 (RUNX2) was down-regulated by bilirubin. This effect was observed at 2 and 24 hours.

With respect to genes involved in osteoclast differentiation and bone resorption, LCA up-regulated the receptor activator of nuclear factor-kappaB ligand (*RANKL*) and the calcitonin receptor (*CALCR)* after 2 hours. This effect was not evident at 24 hours. Similarly, LCA induced a constant overexpression of the matrix Gla protein (*MGP*). No significant effects of bilirubin were observed in the expression of these genes. Conversely, the colony stimulating factor 2 (*CSF2*) (p<0.001) was significantly down-regulated after 2 hours under LCA, but upregulated by bilirubin at 24 hours.

Within the growth factor family, LCA increased Insulin-like growth factor 1 (*IGF1*) expression levels after 2 hours, and bilirubin significantly increased transforming growth factor beta-1 (*TGFB1*) after 24 hours (p=0.03).

LCA significantly up-regulated 1,25-dihydroxyvitamin D3 24-hydroxylase (*CYP24A1)* gene expression, an effect which was observed from the first two hours and sustained throughout of the experiment (p<0.001)

Finally, expression levels of both dentin sialophosphoprotein (*DSPP*) (p<0.01) and decapentaplegic homolog 6 (*SMAD 6*) (p<0.02) were down-regulated after 2 hours under bilirubin treatment, whereas LCA produced the same effect on the Wnt signaling pathway by down-regulating the dickkopf-related protein 1 (*DKK1*) gene expression. This effect was observed after 2 hours of treatment and was much higher at 24 hours (p=0.01).

### Effect of UDCA and interaction with LCA and bilirubin on gene profiles

As compared with controls, UDCA has a significant effect, mainly on three big family genes: apoptosis (figure 3A), bone morphogenetic proteins (BMPs) (figures 2 and 3B) and bone specific genes (osteoblastic transcription factors and specific osteogenic factors) (figure 3C). UDCA resulted in specific gene expression changes by itself or by modifying the effects of LCA and bilirubin. Accordingly, UDCA 10μM and 100μM diminished the gene expression of pro-apoptotic genes *BAX*, *BCL10*, *BCL2L13*, *BCL2L14* and *BCL3*, and counteracted both the LCA and bilirubin effects. UDCA increased the expression of the anti-apoptotic genes *BCL2*, *BCL2A1*, *BCL2L1* and *BCL2L2*, being significant for *BCL2A1* (p=0.03). Moreover, this UDCA anti-apoptotic function neutralized the LCA and bilirubin effects by abolishing the significant decrease produced by them on these anti-apoptotic genes (p<0.03). No significant changes were found in the expression of caspase family genes under UDCA (figure 3A).

**Fig 3.**
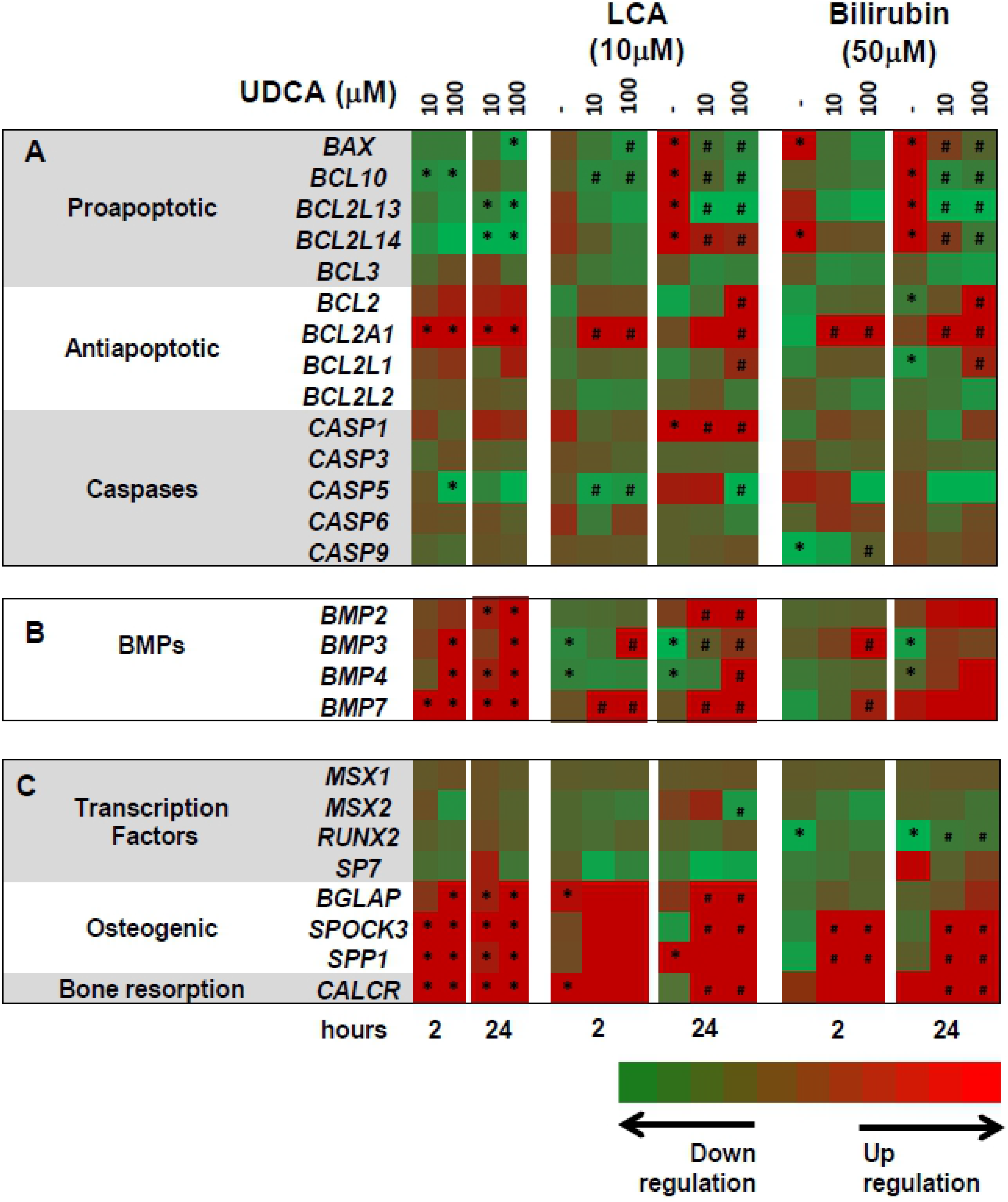
Heat map of the differential gene expression induced by UDCA and neutralizing effects on osteoblastic cells cultured with lithocholic acid (LCA) and bilirubin. A) UDCA apoptotic profile; B) UDCA BMPs profile, and C) UDCA transcription factors and osteogenic markers profiles. The red colour corresponds to genes that are up-regulated, and green colour corresponds to genes that are down-regulated as compared to controls. #indicates significant differences with respect to non-treated cells. In the experiments with lithocholic acid (LCA) or bilirubin # significant differences of UDCA treated cells with respect to LCA or bilirubin treated cells, or with respect to controls.

UDCA significantly increased *BMP2*, *BMP3*, *BMP4* and *BMP7* gene expression from 2 hours of treatment at 10μM and 100μM and neutralized the down-regulation induced by LCA and bilirubin on BMPs expression (p<0.001) (Figures 2 and 3B).

Furthermore, UDCA significantly increased gene expression of the bone resorption marker CALCR and of the osteoblastic specific bone markers *BGLAP*, *SPOCK3* and *SPP1*, when cells were treated with UDCA (10μM or 100μM) or when cells were treated at the same time with 10μM LCA or 50μM bilirubin (Figure 3C).

## Discussion

The effects of substances retained in chronic cholestasis on bone cells have been previously described, suggesting direct harmful effects of bilirubin and LCA on human osteoblastic cells. Janes at al. observed that plasma with high concentrations of bilirubin resulted in decreased human osteoblast-like cells proliferation [19]. Similarly, bilirubin induced a marked dose-dependent inhibition of avian chondrocytes proliferation [20]. Previous studies from our group have shown a decrease of human osteoblastic cells viability and differentiation when bilirubin and sera from jaundiced patients was added to the culture media [7]. Also, an increase in apoptosis was observed in both bilirubin and LCA-treated osteoblastic cells [24]. However, there are scarce data on the targets of these substances in osteoblasts [7,24]. The current study provides new insights in the effects of bilirubin and LCA on the expression of relevant groups of genes, as well as on the involved molecular pathways.

The essential impact of bilirubin on the expression of genes involved in apoptosis in human osteoblasts is confirmed in this study. It should be noted that bilirubin at 50 μM dramatically increases the pro-apoptotic related genes and decreases the anti-apoptotic genes. Both phenomena occur after only few hours of treatment, suggesting that bilirubin could play a major role in the regulation of the apoptotic related genes. This observation leads us to confirm our earlier observations about the pro-apoptotic role of bilirubin on osteoblastic cells [24] and other cell types and tissues [15–16,25,26]. Likewise, the relationship between LCA and apoptosis in different tissues and cell types has been widely demonstrated. Previous studies observed LCA-induced apoptosis mediated by the nuclear receptor Nur77 expression in both human liver and colon cancer cells as well as in mouse hepatocytes [27]. LCA has also a pro-apoptotic effect in human colon adenocarcinoma cell lines [28], in human neuroblastoma cells [29] and in cultured syncytiotrophoblast cells [30]. Finally, LCA induces apoptosis in human osteoblasts, increasing DNA fragmentation, caspase-3 activity and producing an up-regulation and a down-regulation of *BAX* and *BCL2* respectively [24]. Accordingly, the current study describes the apoptotic action of LCA through the increase of pro-apoptotic and the decrease of anti-apoptotic genes.

A new finding described in this study is the relationship between BMPs gene expression under the effects of LCA, bilirubin and UDCA (figures 2 and 3B). BMPs are members of the transforming growth factor-β (TGF-β) superfamily and a subset of BMPs possess the ability to induce bone and cartilage formation and enhance osteogenesis and fracture healing [31–33]. BMPs are extracellular cytokines, originally isolated from bone extract and are produced in nearly all skeletal cells [34,35]. Accordingly, *BMP2* vastly increases osteocalcin, and a short-term expression of *BMP2* is necessary and sufficient to irreversibly induce bone formation [36,37]. Additionally, *BMP7* accelerates calcium mineralization and induces the expression of osteoblastic differentiation markers such as ALP activity [38,39], and loss of both BMP2 and BMP4 results in severe impairment of osteogenesis [40]. The decrease of *BMP3* and *BMP4* expression observed in our study under LCA and bilirubin treatments open up new approaches to illustrate their mechanism of action. On the other hand, the current study clearly demonstrates that UDCA causes an important up-regulation of *BMP2*, *BMP3*, *BMP4* and *BMP7* genes, and in addition counteracts the effects of LCA and bilirubin induced down-regulation of these genes.

The runt-related transcriptional factor 2 (RUNX2) [41] is the key transcription factor involved in osteoblasts differentiation under the guidance of BMPs signaling, since its expression can be induced by both *BMP2* and *BMP7* [42]. The increased expression of BMPs under UDCA treatment leads us to relate it with the expression levels of the transcriptional factor RUNX2, which are down-regulated after bilirubin 50μM administration on cells treated for a short time and maintained steadily for 24 hours. These results are consistent with those published in previous studies, in which a down-regulation of RUNX2 with a consequent decrease in osteoblast differentiation was produced by the cell exposure to 50μM of bilirubin [7] and under low concentrations of bilirubin (3 and 30μM) in rat osteoblasts primary culture with osteogenic medium for 3 or 14 days [43]. The potential beneficial effects of UDCA on bone cells may be partially explained the BMPs up-regulation, which in turn induces RUNX2 expression. These effects were not clearly observed in these experiments, although UDCA partially attenuates the RUNX2 down-regulation induced by bilirubin.

Despite the fact that new insights into gene profiling induced by bilirubin, LCA and UDCA have been described in this study, some limitations should be taken into account. The main concern is that the experiments were carried-out using the human osteosarcoma cell line Saos-2, which although it is most similar cell line to human primary osteoblasts, its behavior could be different. Our next approach will then be to check the gene expression of the most relevant results of our current study in primary human cultures, particularly those related to BMPs and some transcription factors such as RUNX2.

In summary, the current study shows that accumulated products of cholestasis decrease the expression of some BMPs, which are mostly strong osteogenic agents and synergize with osteogenic transcriptional factors such as RUNX2, that was down-expressed as well. Furthermore, the addition of biliary acids and bilirubin up-regulated pro-apoptotic and down-regulated anti-apoptotic genes. These changes in the apoptosis pathways involving the Saos-2 cells could modify the rate of bone formation and, therefore, be considered as a truthful pathogenic mechanism of osteoporosis in cholestatic and in end-stage liver diseases.

